# Post-translational modification fidelity of recombinant human lactopontin expressed in *Kluyveromyces lactis*

**DOI:** 10.64898/2026.05.12.724256

**Authors:** Joshua Excell, Agata Giardina, Emily Sakamoto-Rablah, Kate Royle, David Nunn

## Abstract

Recombinant human lactopontin (rhLPN), an equivalent of human milk lactopontin, is of increasing interest for human nutrition applications due to its roles in mineral binding, gastrointestinal function and immune modulation. These properties depend strongly on post-translational modifications, particularly phosphorylation and glycosylation. Here, we present a comprehensive molecular comparison between rhLPN produced in *Kluyveromyces lactis* at laboratory and pilot scale with native human lactopontin (nhLPN) isolated from human milk. Mass spectrometry-based peptide mapping confirmed the primary structure and identified extensive phosphorylation, consistent with the native protein. Middle-up analyses demonstrated closely matched phosphoform distributions between rhLPN and nhLPN, while glycosylation profiling revealed a defined population of low-complexity O-glycoforms localized to the N-terminus. Functional assessment demonstrated substantially greater iron binding by phosphorylated rhLPN compared with dephosphorylated and non-phosphorylated forms. Similar phosphorylation-dependent behaviour was observed for bovine lactopontin, supporting a conserved role for phosphorylation in mineral interaction. Across five 750 L pilot scale batches, both phosphorylation and glycoform distributions were highly consistent, indicating robust process reproducibility. Together, these results demonstrate that rhLPN produced in *K. lactis* recapitulates key structural and functional attributes of nhLPN, supporting its suitability as a scalable ingredient for nutrition applications.

## 1 Introduction

Osteopontin (OPN), also known as secreted phosphoprotein 1 (SPP1), is a highly multifunctional protein implicated in diverse physiological processes, including bone metabolism and biomineralization, modulation of the immune system, wound healing, intestinal health and brain development (1). A distinct dietary form, lactopontin (LPN), is present in human milk and has been associated with key aspects of infant development, including neurodevelopment (2,3) and gastrointestinal health (4,5). Increasing interest in LPN has extended beyond infancy into adult nutritional applications, such as the promotion of biomineralization in bone remodelling (6).

A defining feature of lactopontin is its extensive post-translational modifications (PTMs), particularly phosphorylation and O-linked glycosylation, which play distinct and critical roles in dictating its nutritional and bioactive properties. Human milk-derived LPN possesses 36 phosphorylation sites per molecule, with approximately 30–32 sites typically occupied (7). This intense clustering of phosphorylation sites is required for coordinating essential minerals such as iron (8) and for mediating binding interactions with other functional, bioactive proteins in milk, including lactoferrin (9). O-glycosylation also provides functional benefit as this modification increases gastrointestinal resistance and allows the functional fragments to survive digestion (10). Importantly, these modifications work in tandem and this cooperative effect is underscored by *in vivo* research which demonstrated that oral administration of natively modified bovine milk OPN significantly attenuated disease parameters in a mouse model of acute colitis, whereas non-modified, unphosphorylated, and non-glycosylated recombinant controls were entirely ineffective (5).

Given these multifunctional roles, there is increasing interest in the use of human lactopontin in nutrition applications. However, sourcing native human LPN (nhLPN) from donor breast milk presents ethical challenges and is neither sustainable nor scalable. Although bovine lactopontin is a commercially available alternative (Lacprodan® OPN-10), species-specific differences in sequence and post-translational modification may limit functional equivalence.

Precision fermentation, therefore, offers a promising route to the production of recombinant human lactopontin (rhLPN), provided that the PTMs can be faithfully reproduced (11,12). Early studies employed *Escherichia coli* (13); however, bacterial systems lack endogenous eukaryotic post-translational machinery, resulting in non-glycosylated proteins that require post-expression phosphorylation *in vitro*. While plant (14) and mammalian platforms (15) have generated rhLPN, the former lacks the extensive phosphorylation which is a hallmark of nhLPN, and the latter has challenges of scalability and cost. Microbial systems such as *Kluyveromyces lactis*, an organism widely used in biotechnology and for the safe manufacture of food ingredients (16), offer scalable, cost-efficient production suitable for nutritional applications. For O-linked glycosylation, yeast hosts typically initiate processing via protein O-mannosyltransferases (PMTs) in the endoplasmic reticulum to yield short, uncharged linear chains restricted to four to five mannose residues, avoiding the hyper-branched complexity seen in higher eukaryotes (17). However, because yeasts lack the endogenous capacity to carry out high-density phosphorylation on secreted dairy targets, a system incorporating a functional Fam20C kinase can be used. Engineering yeast systems to support human-like PTMs in this way was established over ten years ago (18).

In this study, we describe the post-translational modifications of rhLPN produced in *K. lactis* and evaluate its molecular fidelity relative to nhLPN. Using complementary mass spectrometry approaches, we compare phosphorylation and glycosylation profiles at both site-specific and population levels. In addition, we assess functional relevance through iron-binding measurements and evaluate batch-to-batch consistency at pilot scale. This work aims to determine whether recombinant LPN can reproduce both the structural and functional characteristics required for nutrition applications at scale.

## 2 Materials and methods

### 2.1 Materials

Two sources of nhLPN from breast milk have been used. The first is purified lactopontin derived from pooled human breast milk obtained from healthy donors under ethically approved collection protocols. The material represents the naturally occurring form of human lactopontin from milk and is referred to as nhLPN-milk. The second is also purified lactopontin derived from human breast milk, available commercially from Antibodies Online (ABIN1047989, Germany), and is referred to as nhLPN-AB. Additionally, recombinant 6X-His tagged human LPN (residues 17–314) was expressed and purified from *E. coli* using standard protein expression and purification techniques (13), referred to as non-phosphorylated rhLPN. Commercial bovine lactopontin from milk, Lacprodan® OPN-10 (Arla Foods Ingredients Group, Denmark), referred to as nbLPN, was used for comparative iron-binding experiments.

### 2.2 Manufacture of recombinant human lactopontin

Laboratory scale rhLPN was produced in shake flasks using batch fermentation from a strain of *K. lactis*, BD-LPN60, that has been engineered to enable phosphorylation of secreted rhLPN through the co-expression of human Fam20c kinase. The amino acid sequence of rhLPN from BD-LPN60 differs from nhLPN by three positions. Two, I1S and P2V are at positions one and two of the mature protein and have been introduced to facilitate the cleavage of the signal peptide by the *K. lactis* Kex2 secretory protease. The third amino acid change, R232L is in the C-terminal region and has been introduced to improve production of the intact, mature protein. The rhLPN is produced constitutively and secreted into the fermentation broth. At harvest, the fermentation broth is separated from the biomass and filtered, concentrated and the rhLPN purified with an anion-exchange chromatography step. A final dialysis step to remove salt was implemented to enable downstream analytics.

For pilot scale production, a series of five 750L fed-batch fermentations was carried out. For each, a working stock vial of the strain BD-LPN60 was used to inoculate a seed fermenter, which in turn was used to inoculate a production fermenter. The laboratory scale downstream process was implemented at the manufacturing scale, with the addition of a final lyophilization step. Product purity was assessed by LC-MS/MS peptide mapping with Top3-based label-free quantification (19), yielding an average purity of 98.6% (SD = ±0.817%).

### 2.3 Liquid chromatography mass spectrometry (LC-MS)

#### 2.3.1 Peptide mapping with LC-MS

Phosphorylation sites were identified using LC-MS/MS peptide mapping experiments. Samples were digested separately with trypsin, Glu-C and thermolysin. The trypsin and Glu-C digests were performed overnight at 37°C in 20 mM Tris-HCl (pH 7.0) with an enzyme: substrate ratio of 1:20.

The thermolysin digest was performed for 2 hours at 75°C in 20 mM Tris-HCl, 0.5 mM CaCl (pH 8.0) with an enzyme: substrate ratio of 1:10. Both reactions were quenched with 17.5% formic acid and centrifuged to remove particulates. Peptides were analysed on a Q-TOF LC-MS/MS system from Agilent Technologies (6545-XT, USA) using an AdvanceBio Peptide Map 2.1 x 150 mm, 2.7 µm column from Agilent Technologies (653750-902, USA) heated to 60°C. The samples were eluted at a flow rate of 0.4 mL/min by using 1% formic acid in water (solvent A) and 1% formic acid in acetonitrile (solvent B). In 40 min of linear gradient, solvent A decreased from 97% to 60%, while solvent B increased from 3% to 40%. The heated Dual Agilent Jet Stream Electrospray Ionization (Dual AJS ESI) source was operated in positive mode with a gas temperature of 350°C and a flow of 12 L/min. The sheath gas temperature and flow were set to 350°C and 11 L/min, respectively. The analysis was performed using a high-resolution mass spectrometer operated in Extended Dynamic Range (2 GHz) acquisition mode. The source conditions were maintained with a nozzle voltage of 1000V, a fragmentor voltage of 175V, and a skimmer voltage of 65V. Mass calibration was continuously monitored using reference masses at m/z 922.0098 to ensure mass accuracy. Data was acquired over a mass range of m/z 100–3,200 at an acquisition rate of 5 spectra/sec. For the Auto MS/MS experiments, the top 3 precursors per cycle were selected for fragmentation using a narrow isolation width of approximately 1.3 m/z. MS/MS spectra were collected from m/z 50–3,200 at a minimum rate of 3 spectra/sec. The collision energy (CE) was dynamically calculated from the precursor mass according to the following linear equation: CE = 3.6*(m/z)/100–4.8. Raw data acquired from LC-MS/MS were processed using MassHunter BioConfirm B.08.00 software from Agilent Technologies (G6829AA, USA) filtered to remove matches with a score below 85.00. The output of the BioConfirm methodology was validated by comparing the results to data processed by the Find by Formula algorithm in MassHunter Qualitative Analysis software (v10.0) from Agilent Technologies (G3336AA, USA) using a database containing the formulas for digest peptides with all possible levels of phosphorylation occupancy.

#### 2.3.2 Middle-up analysis of thrombin-cleaved material with LC-MS

Phosphoform population profiles were identified with a middle-up analysis of thrombin-cleaved material, which can be resolved by separating N- and C-terminal fragments of the protein, simplifying the signals for deconvolution. Samples in 20 mM Tris-HCl (pH 7) were digested with thrombin from Sigma-Aldrich (T7572, UK) at a ratio of 30 mU/µg LPN for 1 hour at 37°C to generate discrete N-terminal and C-terminal fragments. The resulting fragments were resolved on an Advance Bio RP-mAb C4 column (2.1 x 100 mm, 3.5 µm) from Agilent Technologies (795775-904, USA) using two mobile phases at a flow rate of 0.45 mL/min: 0.1% formic acid in water (solvent A), and 0.1% formic acid in acetonitrile (solvent B). The elution profile began with an initial composition of 95% solvent A and 5% solvent B. A linear gradient was applied over 60 minutes, reaching 50% solvent B, then ramping to 100% solvent B over 1 minute. Finally, the system was re-equilibrated to 5% solvent B. The samples were analyzed using the Dual AJS ESI source operating in positive ion polarity. Desolvation was achieved using a drying gas temperature of 320°C. A capillary voltage of 3500 V and a nozzle voltage of 1000 V were set. MS spectra were acquired over a mass range of m/z 100–3,200. Deconvoluted spectra were used to assess the population of modification states present within each fragment. Deconvolution was run through MassHunter BioConfirm v11.0 software from Agilent Technologies (G6829AA, USA) using the maximum entropy algorithm. The mass range was set to 15000-25000 Da with a mass step of 0.1 Da. The m/z range was set to 600-3000. The mass match tolerance was set to +/− 20 ppm.

#### 2.3.3 Intact analysis of dephosphorylated material with LC-MS

Glycoform population profiles were identified using intact analysis of dephosphorylated material, where removal of phosphorylation enables simpler signal resolution. Samples, in a reaction buffer of 10 mM Tris-HCl, 2 mM MgCl_2_, pH 9.7, were treated with alkaline phosphatase from Sigma-Aldrich (P6774, UK) at a ratio of 1U/10 µg LPN for 5 hours at 37°C to remove phosphate sites. LC separation was performed on an Advance Bio RP-mAb C4 column (2.1 x 100 mm, 3.5 µm) from Agilent Technologies (795775-904, USA) with a flow rate of 0.45 mL/min. The elution profile began with an initial composition of 95% solvent A and 5% solvent B. A linear gradient was applied over 60 minutes, reaching 50% solvent B, then ramping to 100% solvent B over 1 minute. Finally, the system was re-equilibrated to 5% solvent B. The samples were analyzed using the Dual AJS ESI source operating in positive ion polarity. Desolvation was achieved using a drying gas temperature of 320°C. A capillary voltage of 3500 V and a nozzle voltage of 1000 V were set. MS spectra were acquired over a mass range of m/z 100–3,200. Data was analysed with Bioconfirm software.

#### 2.3.4 MALDI-TOF analysis of glycan structures

Glycan structures were identified following release using the O-FANGS approach (20). O-glycans were released using a 16-hour on-filter incubation in 25% ammonium hydroxide at 45°C. Released, dried, free O-glycans were permethylated with iodomethane. MALDI-MS spectra were acquired in positive ion, reflectron mode using a Bruker UltrafleXtreme mass spectrometer. Glycan assignment was performed by comparing measured m/z values with theoretical glycan masses in the GlyConnect database via the GlycoMod interface, using a mass tolerance of 0.1 Da.

### 2.4 Measurement of iron binding

The iron binding capacity of rhLPN, dephosphorylated rhLPN, native bovine LPN (nbLPN), dephosphorylated nbLPN and non-phosphorylated rhLPN was quantified colourimetrically using 3-(2-Pyridyl)-5,6-di(2-furyl)-1,2,4-triazine-5′,5′′-disulfonic acid disodium salt (Ferene-S) from Sigma-Aldrich (82940, UK), which forms a blue complex with ferrous iron (8). Dephosphorylated rhLPN and nbLPN were produced as described in Section 2.3.3.

For each sample, 100 µL of protein at 0.1 mg/mL in 50 mM MES buffer (pH 6.7) was incubated with FeSO_4_.7H_2_O at various concentrations for 30 minutes at room temperature. Unbound iron was removed by desalting with a Zeba Spin desalting plate from Thermo Fisher Scientific (89808, USA). The sample was freeze-dried, re-suspended in 50 µL of 65% nitric acid, and incubated at 70°C for 2 hours. After cooling to room temperature, the samples were neutralized with 70 µL of 10 M NaOH, and 750 µL of 5 mM Ferene-S, 0.2 M L-ascorbic acid in 0.4 M ammonium acetate buffer (pH 4.3) was added. Color development was measured after 30 minutes by absorbance at 595 nm. A standard curve of FeSO_4_ in water was used to interpolate bound iron. The presence of iron in control samples containing no protein was found to be independent of Fe concentration, with a mean value of 0.80 ug/ml, which was background-subtracted from all subsequent measurements. Iron binding values measured for 0.01 U/µl alkaline phosphatase were subtracted from dephosphorylated rhLPN data to account for the presence of alkaline phosphatase in those samples. All measurements were performed in triplicate. Iron-binding data were fitted using an asymmetric five-parameter nonlinear regression model to account for the heterogeneous binding behaviour expected from multiple phosphorylated binding regions.

### 2.5 Statistical analysis

#### 2.5.1 Analysis of phosphorylation-state distributions

Relative abundance data (% quantitative peak area) for each phosphorylation state were obtained from three replicate analyses per batch. Multivariate differences in phosphorylation-state distributions between batches were assessed using permutational multivariate analysis of variance (PERMANOVA), implemented in the *vegan* package in R. Analyses were performed on replicate-level data using Bray–Curtis dissimilarities with 9,999 permutations. Homogeneity of multivariate dispersion was evaluated using the *betadisper* function to confirm that observed differences were attributable to shifts in distribution centroids rather than differences in within-batch variance (Note: Negative eigenvalues arising from non-Euclidean distance structure were handled using standard correction within the *betadisper* implementation). Principal component analysis (PCA) was applied to visualize similarities and differences between batches and to identify the phosphorylation states that contributed most to variance among samples. PCA results were visualized using biplots displaying both batch scores and phosphorylation-state loadings.

To visualize phosphorylation-state distributions, mean relative abundances across replicate analyses were calculated for each batch and displayed as heat maps.

#### 2.5.2 Analysis of glycoform distributions

Relative abundances of glycoforms were determined for each batch. Glycoform abundance data were treated as compositional and therefore subjected to centered log-ratio (CLR) transformation prior to principal component analysis. PCA was then applied to the transformed data to examine similarities and differences in glycoform profiles between batches. PCA results were visualized using biplots that display batch scores alongside glycoform loadings. Formal multivariate hypothesis testing was not performed on glycoform data since only a single analytical measurement was available for each batch.

Heat maps were used to visualize glycoform distributions across batches. Glycoforms were ordered using hierarchical clustering based on Euclidean distances to group species with similar batch-to-batch abundance patterns.

## 3 Results

### 3.1 Confirmation of rhLPN phosphorylation sites and comparison to native human LPN from milk

Peptide mapping by LC-MS/MS was performed for two purposes; first to confirm the expected primary structure of rhLPN, and second to localise sites of phosphorylation within the protein sequence. The phosphorylation sites of two sources of native human lactopontin, nhLPN-milk and nhLPN-AB, were also mapped. These three datasets were compared with previously reported patterns for milk-derived lactopontin (7), confirming that rhLPN produced in *K. lactis* possesses a phosphorylation landscape consistent with that of nhLPN (Figure 1).

**Figure 1.**
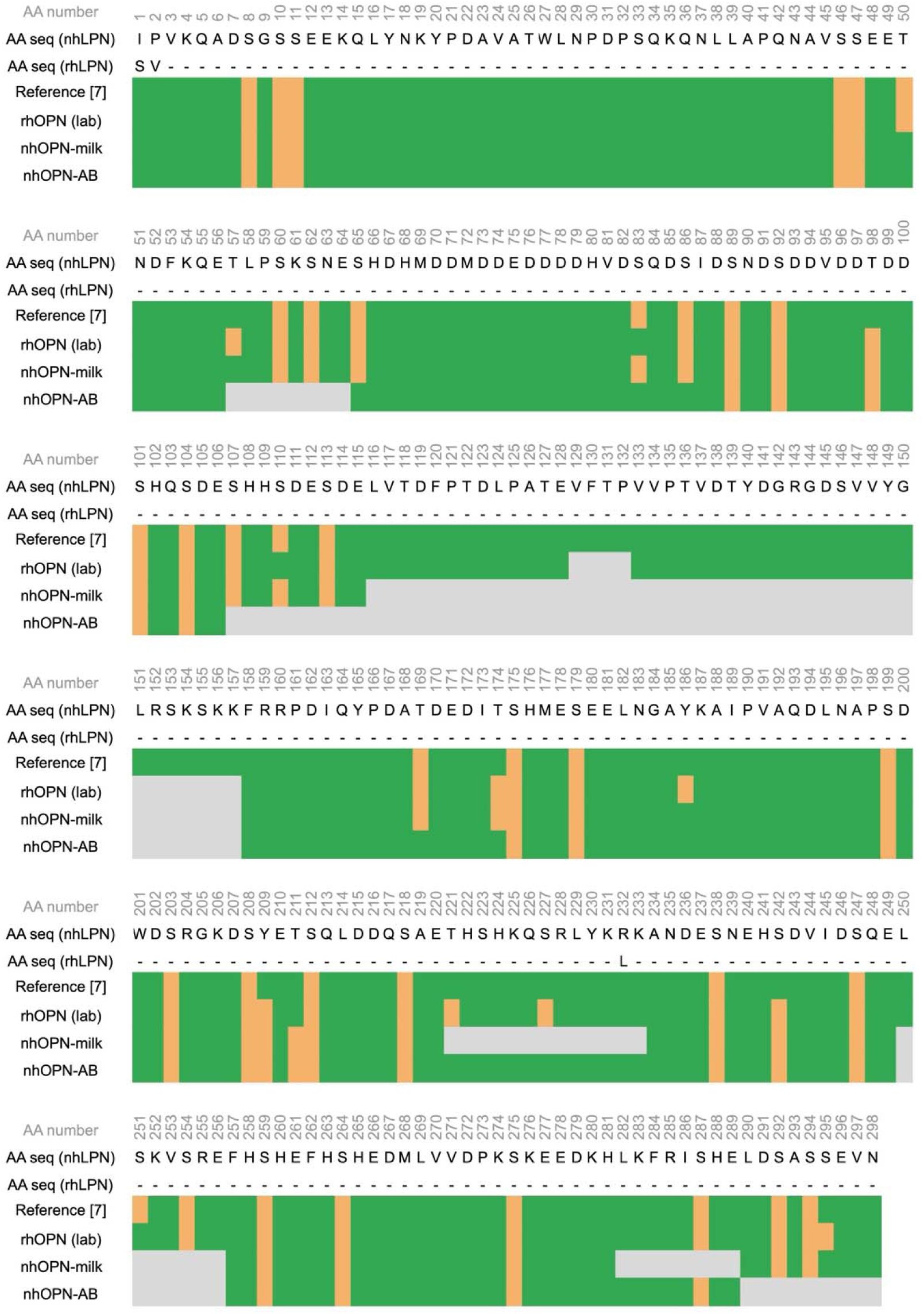
Phosphorylation site profiles of rhLPN and nhLPN. The 298 amino acid sequence for each protein, without the signal peptide, is shown. Dashes indicate homology; the three single amino acid substitutions in rhLPN are described in Section 2.2. Green bars indicate peptide sequence coverage, where grey bars indicate the absence of peptide sequence coverage. Orange bars indicate phosphorylation sites. Sequence coverage for each of the samples was as follows: rhLPN (lab): 96.3%, nhLPN-milk: 76.5%, and nhLPN-AB: 74.8%.

Of the 36 published sites, 33 were detected in rhLPN (Ser8, Ser10, Ser11, Ser46, Ser47, Thr50, Ser60, Ser62, Ser65, Ser86, Ser89, Ser92, Ser101, Ser104, Ser107, Ser113, Thr169, Ser175, Ser179, Ser199, Ser203, Ser208, Ser212, Ser218, Ser238, Ser247, Ser254, Ser259, Ser264, Ser275, Ser287, Ser292 and Ser294, where numbering begins at Ile1 for the mature, secreted human LPN protein). The remaining three sites, which have previously been reported but which were not detected, were Ser83, Ser110, and Ser251.

Nine sites not previously identified were detected in rhLPN (Thr57, Thr98, Thr174, Tyr186, Tyr209, Thr221, Ser227, Ser242, Ser295); two of these are tyrosine residues (Tyr186, Tyr209). Of these nine sites, three (Thr98, Tyr209, Ser242) were detected in both sources of nhLPN, and one (Thr174) was identified in one, nhLPN-milk. Two sites (Thr221 and Ser227) were identified in regions where nhLPN sequence coverage was insufficient, preventing conclusions regarding their presence or absence in native LPN.

Given the very similar mass shifts associated with phosphorylation and sulphation in peptide mapping experiments (+79.9663 Da and +79.9568 Da, respectively), phosphorylation-site assignments were further evaluated following treatment of rhLPN and nhLPN-milk with alkaline phosphatase prior to repeat LC-MS/MS analysis. Following treatment, modification masses corresponding to these sites were no longer detected in either sample, supporting their assignment as phosphorylation events. Although alkaline phosphatase exhibits a very strong kinetic preference for phosphate monoesters relative to sulfate monoesters, weak sulphatase activity has been reported and therefore sulphation cannot be excluded unequivocally on the basis of enzyme treatment alone (21). Nevertheless, the assignments are further supported by the prior identification of Tyr209 phosphorylation in recombinant human lactopontin produced in HEK293 cells (15). In addition, sulphotyrosine formation is not generally associated with yeast expression systems (22,23), providing further support for phosphorylation-based assignment of the corresponding rhLPN modifications.

There were 21 consistent sites between previously reported patterns and the three sources of LPN, rhLPN, nhLPN-milk, and nhLPN-AB (Ser8, Ser10, Ser11, Ser46, Ser47, Ser89, Ser92, Ser101, Ser104, Ser175, Ser179, Ser199, Ser203, Ser208, Ser212, Ser218, Ser238, Ser247, Ser259, Ser264, and Ser275). An additional seven sites were observed between previously reported patterns and one of the two LPN sources, due to the absence of sequence coverage (Ser60, Ser62, Ser107, Ser113, Ser287, Ser292, and Ser294). This observation of 28 sites overall supports similarity in phosphorylation localization.

Minor differences were, however, observed. As noted above, differences were observed between published data and rhLPN, as well as between published data and nhLPN. There were seven reported sites which were not detected in one, or both, of the nhLPN sources (Thr50 - both, Ser65 - one, Ser83 - one, Ser86 - one, Thr169 - one, Ser242 - both and Ser287 - one), and four sites in one or both of the nhLPN sources that had not been previously reported (Thr98 - both, Thr174 - one, Tyr209 - both, Thr211 - one). Even between the two sources of nhLPN, six sites present in nhLPN-milk were not detected in nhLPN-AB (Thr50, Ser83, Ser86, Thr169, Ser179, and Ser212). This indicates that, while the overall phosphosite set is broadly conserved, a small subset of sites is variable in native human LPN. While based on a limited number of samples, these observations align with previous reports indicating heterogeneity in lactopontin (24) and dairy protein phosphorylation (25), across milk sources and lactation phases.

### 3.2 Evaluation of the population of phosphoforms

In addition to mapping the phosphorylation sites of human LPN, it was important to evaluate their site occupancy within the purified protein population. As the deconvolution of mass-to-charge data of intact phosphorylated and glycosylated lactopontin was not practical, the proteins were digested with thrombin to separate the phosphorylated C-terminus from the phosphorylated and glycosylated N-terminus.

There are 18 identified sites in the C-terminus of human LPN. Intact mass analysis by LC-MS of the C-terminus of human lactopontin showed a population of phosphorylation isoforms, with each isoform separated by 80 Da, equivalent to the addition of a phosphate group (Figure 2). The most prevalent rhLPN phosphoform contained 14 of the 18 identified C-terminal phosphorylation sites, compared with 12 of 18 sites in the most prevalent milk-derived nhLPN form. Human lactopontin has been reported to contain up to 36 potential phosphorylation sites, with approximately 30–32 phosphate sites per molecule in milk (7). While phosphorylation is not expected to be uniformly distributed across the protein, the C-terminal region accounts for half of the potential sites. Extrapolation of C-terminal occupancy, assuming similar average occupancy across both regions, therefore, suggests overall phosphorylation levels of approximately 28 sites for rhLPN and 24 sites for nhLPN, values that are broadly consistent with published estimates for native lactopontin.

**Figure 2.**
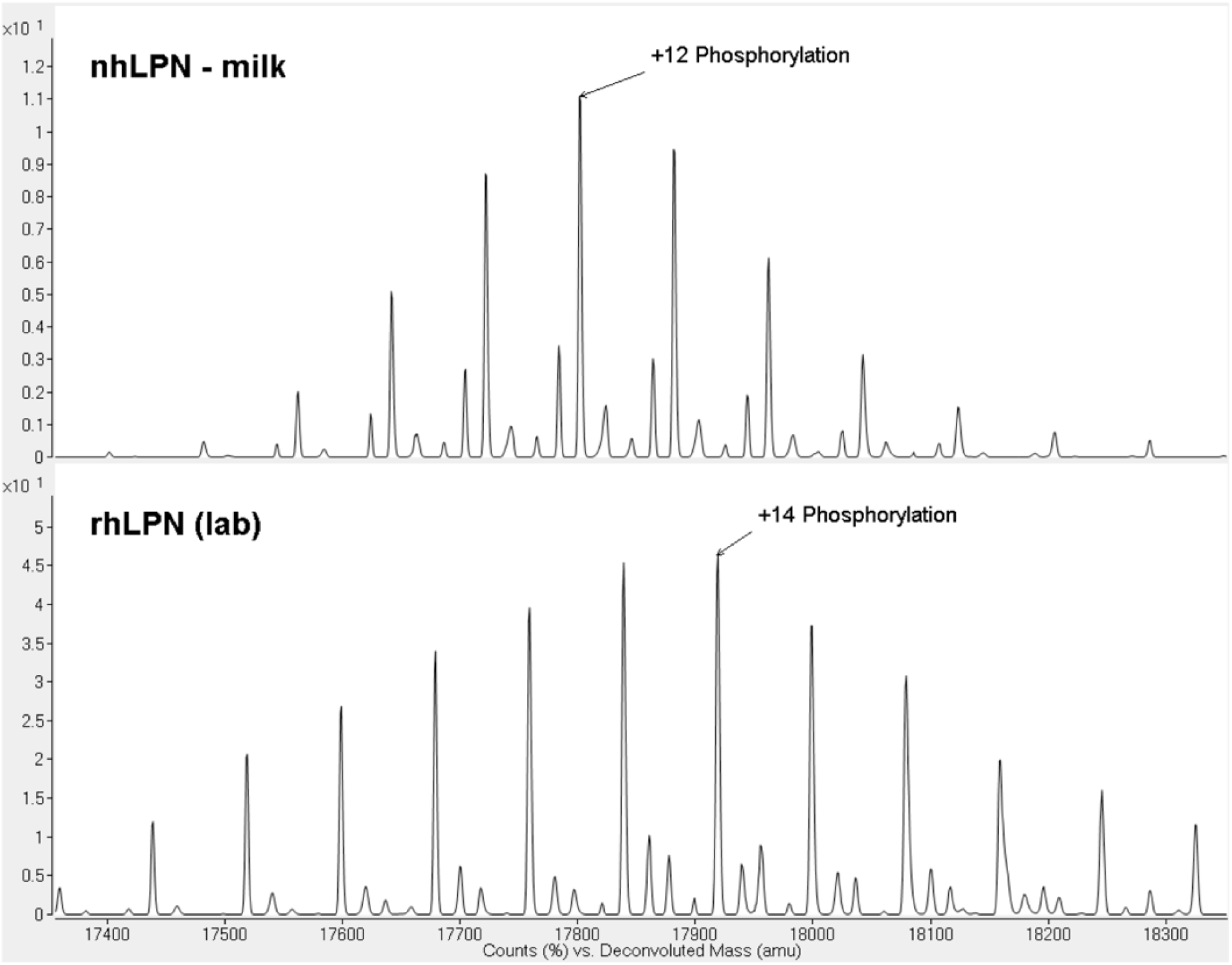
Phosphoform population analysis of the C-terminal region of human lactopontin. Intact LC-MS analysis of the C-terminal fragment following thrombin cleavage reveals a population of phosphorylation isoforms separated by 80 Da increments, characteristic of phosphate group addition. The dominant rhLPN phosphoform contains 14 phosphate groups, compared to 12 in the nhLPN-milk sample. Peaks were identified by adding incremental phosphorylation masses to the expected mass of the unmodified C-terminal fragment. These calculated values were compared against the deconvoluted experimental data, with structural assignments made based on a mass error tolerance limit of 30 ppm. The minor shift in baseline molecular weight between rhLPN and nhLPN-milk is due to three amino acid substitutions in rhLPN, described in Section 2.2 and Figure 1.

### 3.3 Characterization of rhLPN O-linked glycan structures and comparison to native human LPN from milk

During production, recombinant proteins can undergo N-linked and O-linked glycosylation by *K. lactis*. Human lactopontin is a glycoprotein with two unoccupied N-linked glycosylation sites, and five O-linked glycosylation sites (7). To evaluate the glycosylation profile of rhLPN, thrombin cleavage was performed on dephosphorylated laboratory scale material, and the resulting fragments were analyzed by intact LC-MS (Figure 3). The two products of unmodified human LPN, cleaved by thrombin, have expected molecular weights of 16.865 kDa (N-terminus) and 16.799 kDa (C-terminus). In human LPN, the five O-glycosylation sites are Thr118, Thr122, Thr127, Thr131 and Thr136, and are N-terminal to the thrombin cleavage site at Arg168–Ser169. A single peak at 16,799.65 Da indicates that the glycosylation is localized to the N-terminal region of the protein, as expected, as this mass aligns with the mass of the unmodified C-terminal fragment.

**Figure 3.**
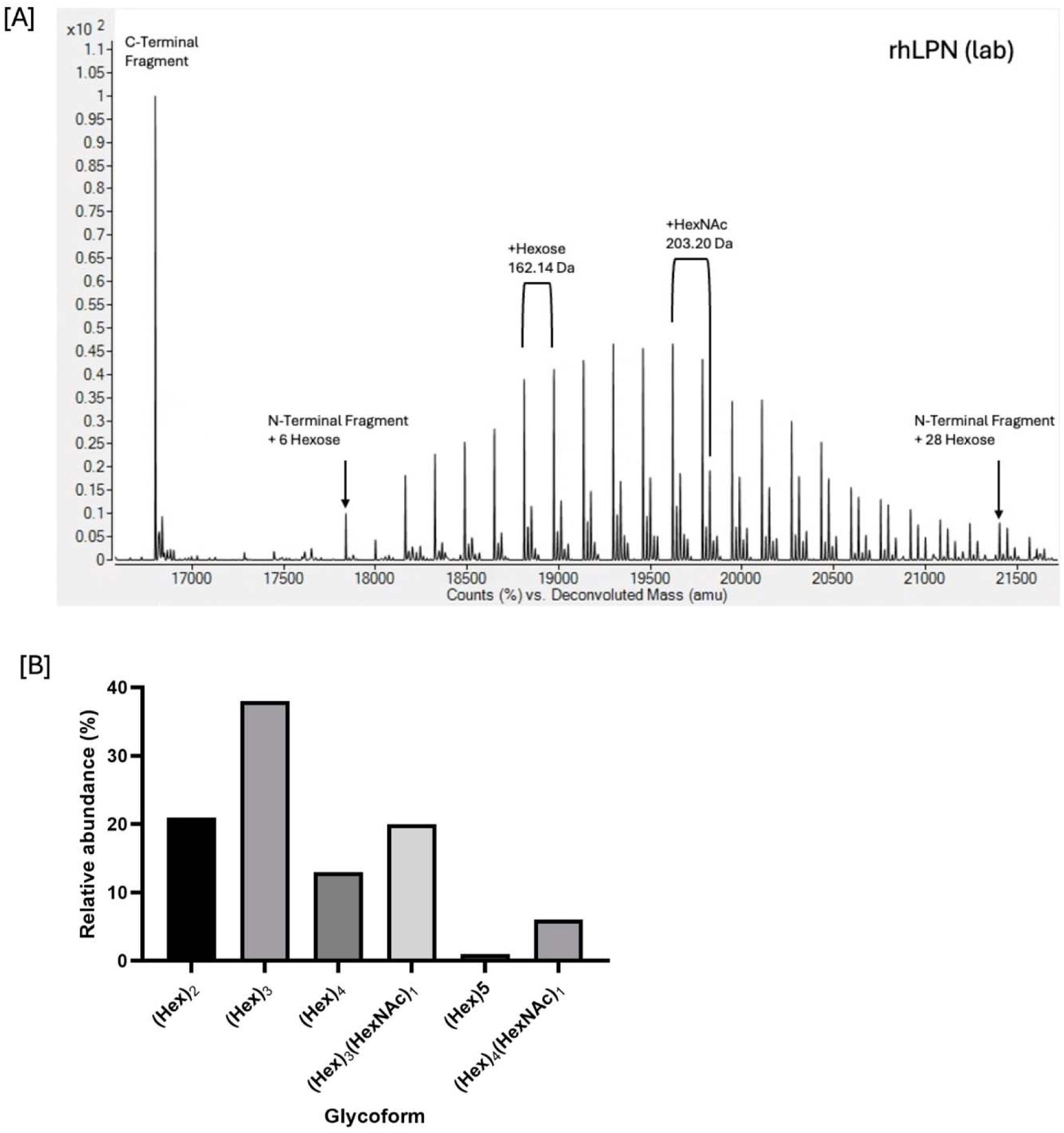
Glycoform analysis of rhLPN. [A] Intact LC-MS analysis of the N- and C-terminal regions of rhLPN following dephosphorylation and thrombin cleavage, showing a largely unmodified C-terminal fragment and a population of N-terminal glycoforms ranging from 6–28 hexose additions. Expected mass differences for the addition of hexose (Hex) and N-acetylhexosamine (HexNAc) are indicated, aligning with the structural data presented in [B]. Deconvoluted experimental masses were compared against the calculated theoretical mass of the unmodified N-terminal fragment (16,865.5257 Da). [B] Structural analysis of O-linked glycans chemically released from the intact protein and analysed by MALDI-TOF mass spectrometry. Detected structures are expressed as a percentage of the total glycan population.

Analysis of the N-terminal population revealed a heterogeneous distribution of glycoforms, ranging from 17838.65 Da to 21567.74 Da, corresponding to species containing 6-29 hexose residues. This distribution is consistent with glycosylation at five sites, with individual glycans comprising approximately one to five monosaccharide units, similar to nhLPN from milk (7). Within this distribution, minor peaks corresponding to HexNAc-modified glycoforms (e.g. (Hex)_3_(HexNAc)_1_ and (Hex)_4_(HexNAc)_1_) were also observed, indicating a low level of N-acetylhexosamine incorporation as previously reported for *K. lactis* (26,27).

To confirm these potential structures, O-linked glycans were chemically released from rhLPN and analysed by MALDI-TOF mass spectrometry (Figure 3 [Inset]). A population of relatively simple glycan structures were detected, with the highest abundance structures being smaller (Hex)_3_ (42% ± 5%) and (Hex)_2_ (24% ± 4%) glycans, with larger (Hex)_4_ (12% ± 2%) and (Hex)_5_ (1% ± 1%) structures present at lower abundances. HexNAc containing structures were detected at a lower relative abundance than structures which were composed of hexose alone ((Hex)_3_(HexNAc)1:17% ± 5%; (Hex)_4(_HexNAc)_1_: 4% ± 2%)). The hexose sugars in these yeast O-glycans are likely to be mannoses and so differ from mammalian glycans where N-acetylgalactosamine, galactose, N-acetylglucosamine, fucose, and sialic acid are typically found (28). Both rhLPN and nhLPN share glycoforms that are comparable in size and complexity, with one to five monosaccharide units, though they differ in sugar composition as expected for a yeast host.

### 3.4 Interactions with the milk matrix: mineral binding

One function of the extensive phosphorylation of nhLPN is to confer a high negative charge density that enables interactions with other bioactive components of the milk matrix, including minerals and bioactive proteins such as lactoferrin. We therefore evaluated the contribution of phosphorylation to iron binding using recombinant human lactopontin (rhLPN) alongside milk-derived, native bovine LPN (nbLPN) preparations differing in phosphorylation status (Figure 4). To benchmark the observed iron-binding behaviour against a previously characterized system, nbLPN preparations were included as comparative controls, as phosphorylation-dependent iron binding has previously been reported for nbLPN (8).

**Figure 4.**
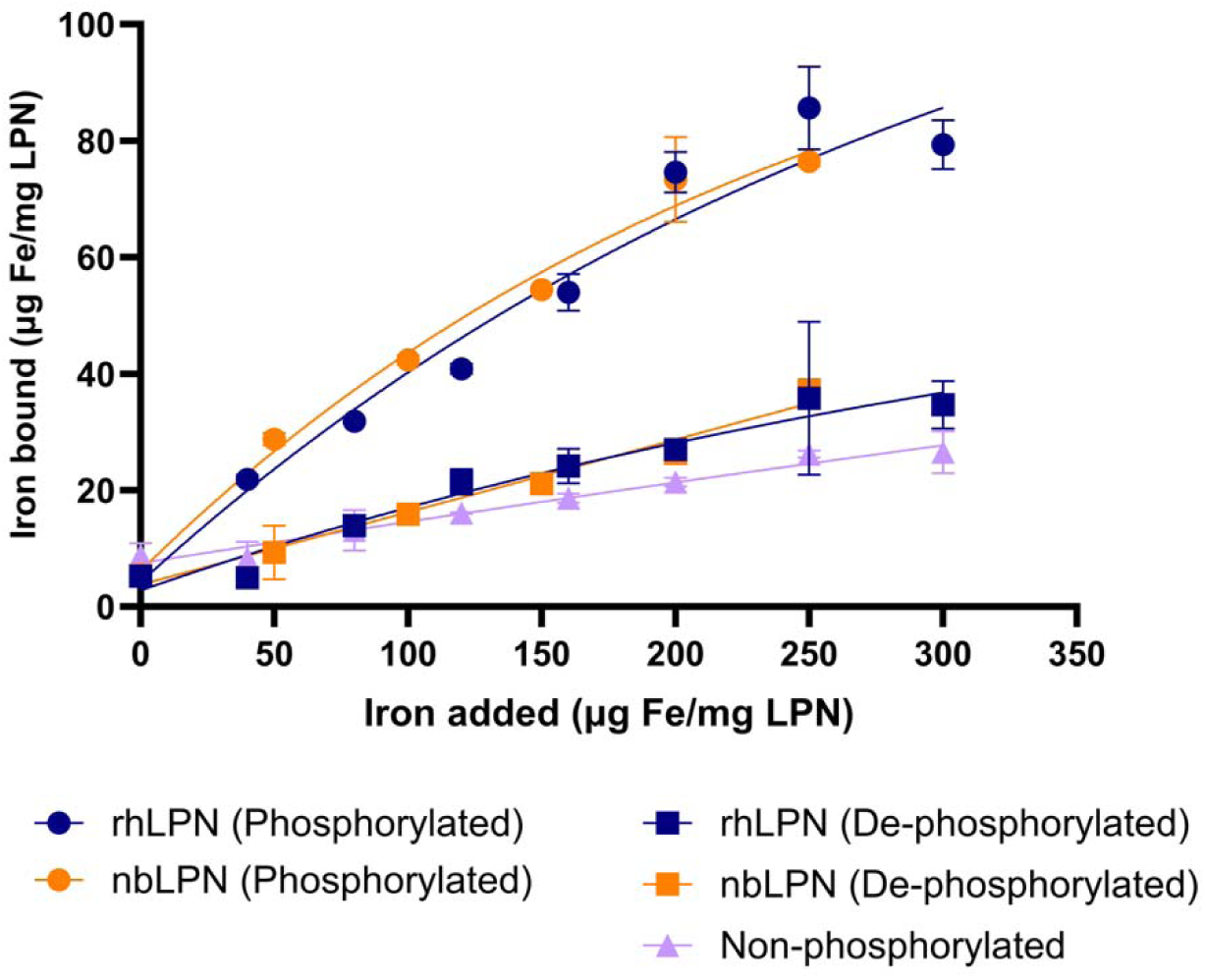
Iron binding profiles of recombinant human and bovine lactopontin phosphorylation variants. Iron binding was measured colourimetrically using the Ferene-S assay following incubation with increasing concentrations of FeSO_4_. Mean values from three replicates are shown for phosphorylated rhLPN, dephosphorylated rhLPN, non-phosphorylated rhLPN expressed in *E. coli*, phosphorylated bovine LPN, and dephosphorylated bovine LPN. Error bars represent the standard error of the mean. Phosphorylated forms of both human and bovine lactopontin exhibited substantially greater iron binding than their corresponding dephosphorylated or non-phosphorylated counterparts, consistent with phosphorylation-dependent mineral interaction.

Consistent with those observations, phosphorylated forms of both human and bovine milk lactopontin exhibited substantially greater iron binding than their corresponding dephosphorylated or non-phosphorylated forms. In contrast, dephosphorylated rhLPN and nbLPN showed markedly reduced iron association across the tested iron concentrations, while non-phosphorylated rhLPN expressed in *E. coli* exhibited minimal binding.

The similar phosphorylation-dependent trends observed for both recombinant human and bovine LPN support the conclusion that phosphate-mediated mineral interaction is conserved across lactopontin species. Together, these findings demonstrate that the phosphorylation introduced during recombinant production of rhLPN is functionally active and recapitulates a key biochemical property associated with native milk-derived lactopontin.

### 3.5 Robustness of the post-translational modification profile across pilot scale batches

To evaluate the consistency of PTM profiles across 750L pilot scale production batches, phosphorylation-state and glycoform distributions were assessed using heat map visualization and multivariate statistical analysis. Transitioning from a laboratory scale environment to a 750 L pilot bioreactor inherently introduces significant shifts in bioprocess dynamics. While the laboratory scale process relied on passive batch dynamics within shake flasks with fixed chemical buffering and depleting nutrients, the pilot-scale strategy utilized a mechanically agitated and sparged bioreactor. This larger-scale system employed automated, closed-loop feedback controls to rigidly maintain fermentation parameters such as dissolved oxygen and pH at setpoints, coupled with a regulated fed-batch feeding strategy to sustain a controlled metabolic steady state.

#### 3.5.1 Batch to batch consistency of phosphorylation-state distributions

To evaluate the phosphorylation-state population, relative abundance data (% quantitative peak area) for each phosphorylation state were obtained from three replicate analyses per batch (Figure 5). Analysis of pilot scale material demonstrated that recombinant human lactopontin (rhLPN) comprised a heterogeneous mixture of mid- and highly phosphorylated species across all batches (Figure 6[A]). The most abundant phosphoforms, P16 and P17, were consistently observed in all batches, representing a modest shift toward higher phosphorylation states compared with laboratory scale material, where P14 was predominant. This shift likely reflects the enhanced metabolic stability and optimised kinetics of the co-expressed human Fam20C kinase within the highly controlled fed-batch bioreactor environment, relative to the drifting, nutrient-depleting conditions of the passive laboratory flasks, although no direct assessment of kinase activity was performed.

**Figure 5.**
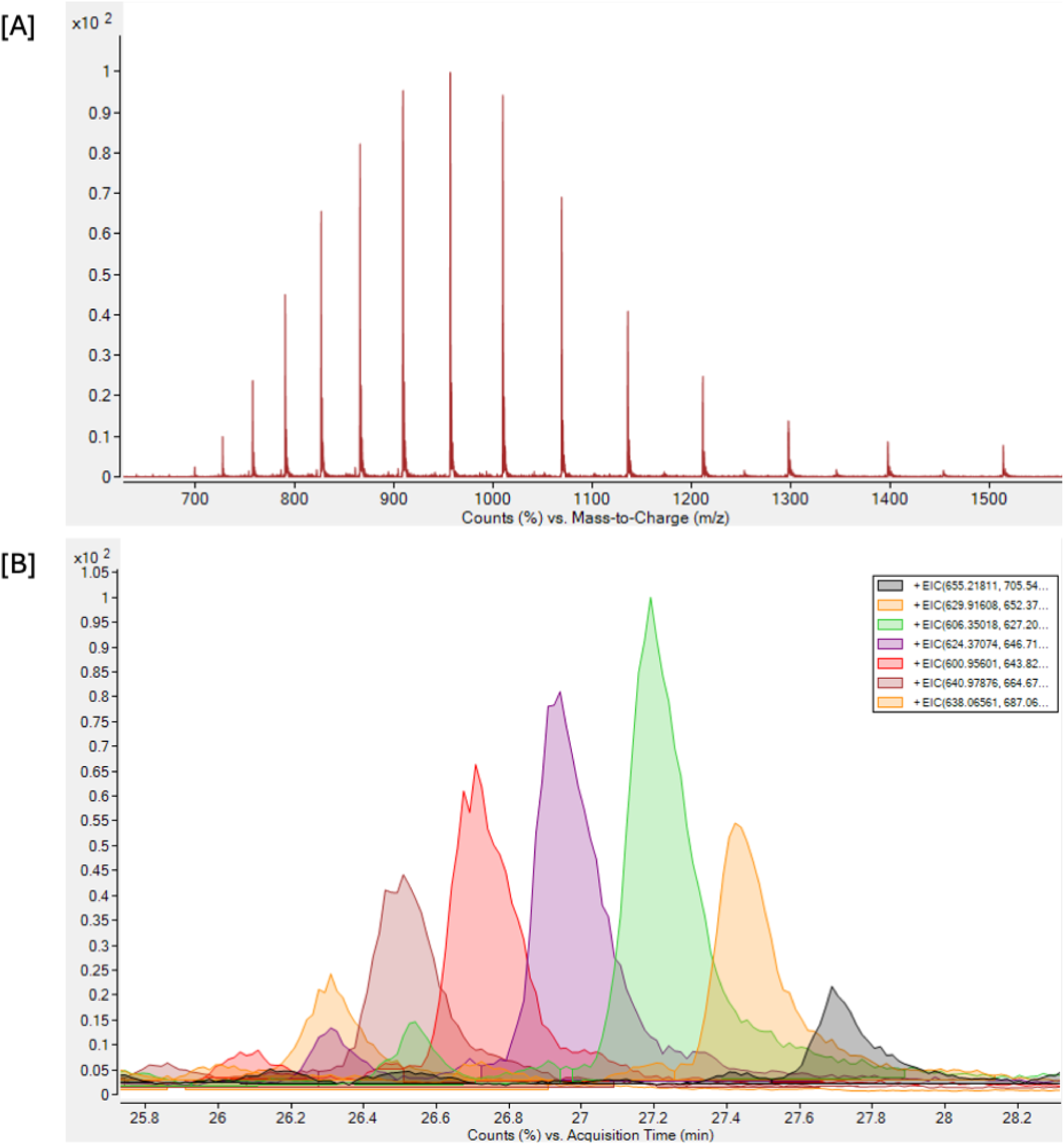
Deconvolution and integration of middle-up data for PCA analysis. [A] The charge state distribution identified for a single phosphorylated C-terminal fragment of production scale rhLPN following thrombin digest. These charge states were deconvoluted using the maximum entropy algorithm, as described in the methods section. [B] An overlaid extracted ion chromatogram (EIC) showing the distribution of deconvoluted masses corresponding to the C-terminal fragment of rhLPN with varying degrees of phosphorylation. Each colour within the legend, and its corresponding peak, represents a deconvoluted mass.

**Figure 6.**
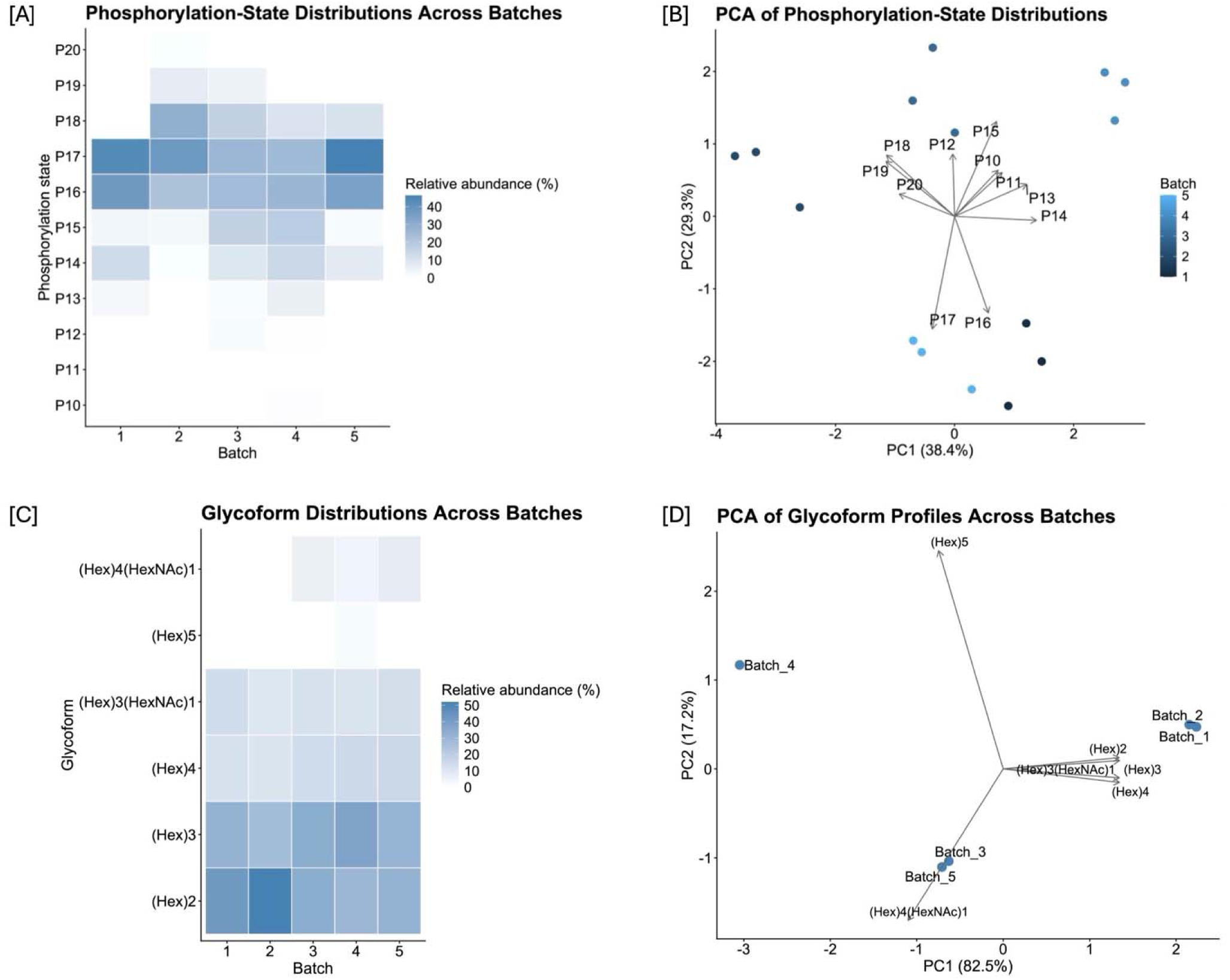
Phosphoform and glycoform distributions across production batches of recombinant human lactopontin. [A] Heat map of mean relative abundances (% quantitative peak area, normalized within each batch) of phosphorylation states across five production batches (n = 3 technical replicates per batch). [B] Principal component analysis (PCA) of replicate-level phosphorylation-state profiles based on relative abundance vectors. Technical replicates cluster tightly, while batch-level separation is limited, indicating minor variation in phosphorylation-state distributions across batches. [C] Heat map of relative abundances (% quantitative peak area) of detected O-linked glycoforms across the same batches. [D] PCA of glycoform abundance profiles. As only a single analytical measurement was obtained per batch for glycoform analysis, PCA is presented for qualitative assessment of global similarity rather than statistical inference.

Relative contributions of individual phosphorylation states varied slightly between batches, indicating minor redistribution within the overall phosphoform population. Principal component analysis (PCA) of replicate-level phosphorylation-state profiles showed tight clustering of technical replicates within each batch, demonstrating high analytical reproducibility (Figure 6[B]). Limited separation between batches was observed along the first principal component, which accounted for a modest proportion of the total variance. Consistent with this, permutational multivariate analysis of variance (PERMANOVA) indicated no statistically significant differences in phosphorylation-state distributions between batches (Bray–Curtis dissimilarity, 9,999 permutations; *R²* = 0.0286, *F* = 0.383, *p* = 0.724). Analysis of multivariate dispersion confirmed comparable within-batch variability (*betadisper* ANOVA, *p* = 0.894), indicating that differences in dispersion did not confound the PERMANOVA result.

Inspection of PCA loadings indicated opposing contributions from lower- to mid-phosphorylation states (P13–P16) and higher phosphorylation states (P18–P20), suggesting that the limited batch-to-batch variation observed was primarily driven by modest redistribution between these phosphorylation populations rather than by the emergence or loss of specific phosphoforms. The absence of statistically significant differences despite visible clustering in PCA reflects the relatively small magnitude of batch-to-batch variation compared with within-population heterogeneity.

#### 3.5.2 Glycoform distributions across production batches

Glycoform profiling revealed a consistent distribution of predominantly low-complexity O-linked glycans across all production batches (Figure 6[C]). The most abundant glycoforms corresponded to (Hex)_2_ (38 ± 9%) and (Hex)_3_ (32 ± 4%), with (Hex)_4_ structures having a smaller contribution (14 ± 2%). HexNAc-containing derivatives were also detected at lower relative abundance ((Hex)_3_(HexNAc)_1_: 12 ± 2%; (Hex)_4_(HexNAc)_1_: 4 ± 4%)). Heat map visualization demonstrated broadly similar glycosylation patterns across batches, with no evidence for the emergence of batch-specific glycoform species.

Principal component analysis of centered log-ratio (CLR)-transformed glycoform abundance data was performed to assess global similarities in glycosylation profiles (Figure 6[D]). As only a single analytical measurement was available per batch, this analysis was interpreted qualitatively. The PCA biplot indicated close clustering of batches in multivariate space, suggesting a high degree of similarity in glycoform distributions across the production runs. Hierarchical clustering of glycoforms further supported this observation, grouping structurally related species with similar abundance patterns across batches.

#### 3.5.3 Overall assessment of PTM consistency

Taken together, these analyses demonstrate that both phosphorylation-state and glycoform profiles of rhLPN are largely conserved across pilot scale production batches. Observed variability is limited to modest redistribution within heterogeneous PTM populations rather than the introduction of new modification states or substantial shifts in overall composition. Such variation is consistent with the known heterogeneity of native lactopontin populations; evaluating what influence this may have on digestion and absorption is the focus of future studies.

## 4 Discussion

This study provides a detailed molecular comparison of recombinant human lactopontin (rhLPN) produced at laboratory and pilot scale in *K. lactis* with native human lactopontin isolated from milk (nhLPN). Comprehensive mass spectrometry analyses confirmed the correct primary structure of rhLPN and revealed closely matched phosphorylation-site occupancy and phosphoform distributions relative to nhLPN. Although differences in the identity of sugar residues within the O-linked glycans were anticipated due to the distinct host cell systems, the resulting glycan structures remained comparatively small and simple in both proteins. While alternative microbial expressions of lactopontin have been reported in platforms like *Komagataella phaffii* (29) and transgenic microalgae (30), these studies focused primarily on secretory titers or purification options without resolving site-specific PTM distributions. By systematically characterising these profiles here, and comparing it to nhLPN, this work addresses a gap in the structural understanding of rhLPN produced from a yeast system.

Importantly, functional analysis supported the biological relevance of the observed phosphorylation landscape. Phosphorylated rhLPN exhibited substantially greater iron binding than dephosphorylated or non-phosphorylated controls, consistent with the established role of clustered phosphate groups in mineral binding and coordination (7). Inclusion of bovine LPN as a comparative control further strengthened the functional interpretation of these experiments, as phosphorylation-dependent mineral interaction has previously been established for bovine milk lactopontin (8). The observation of similar behaviour in rhLPN indicates that the phosphorylation introduced during recombinant production recapitulates a conserved functional property of milk-derived lactopontin. These findings indicate that the phosphorylation introduced during recombinant production is not only structurally similar to that of nhLPN, but also functionally active.

While some differences in phosphorylation at individual sites were observed between rhLPN, nhLPN, and previously published datasets, similar variability was also evident among independent nhLPN sources and reported native datasets. This suggests that modest site-specific variation can occur within an otherwise well-conserved phosphorylation landscape, such that functional equivalence is more appropriately evaluated at the level of overall phosphoform populations and emergent biochemical properties rather than strict conservation of every individual modification site. In this context, the present study demonstrates that rhLPN produced in *K. lactis* successfully recapitulates the population-level phosphorylation characteristics of the native protein while maintaining tight batch-to-batch consistency during pilot-scale production. Collectively, these data support the suitability of *K. lactis*-derived rhLPN as a scalable and functionally relevant alternative for human nutrition applications.

Future studies will focus on further functional validation of *K. lactis*-derived rhLPN, including assessment of its behaviour under simulated gastrointestinal digestion and *in vitro* absorption models. In parallel, *ex vivo* studies will be undertaken to improve understanding of how dietary rhLPN interacts with the immune system, alongside mechanistic investigations aimed at clarifying the role of this key bioactive protein in human nutrition.

## 5 Data Availability Statement

The original contributions presented in this study are included in the article, further inquiries can be directed to the corresponding author.

## 6 Ethics Statement

Human breast milk used in this study was obtained from Prolacta Bioscience (USA). The milk was pooled from healthy donors and collected under supplier managed, ethically approved protocols with written informed consent from all donors, in accordance with applicable regulatory and ethical standards as stated by the supplier.

The material provided was fully de-identified, and this study did not involve direct interaction with human participants. As such, this work does not constitute human subjects research and did not require ethical review or approval.

## 7 Author Contributions

JE, AG and ES-R designed and performed the experiments. JE, AG, ES-R and KR carried out data analysis, statistical analysis and produced the figures. JE and AG contributed to the development of the mass spectrometry methods. KR supervised the study and DN provided strategic oversight and guidance for the project. KR wrote the manuscript with contributions from JE, AG and ES-R. All authors read and approved the final manuscript.

## 8 Acknowledgements

The authors sincerely thank Holly Abell for supporting the experiments and thoughtful discussions. The authors also thank Camila Cotrim for the method development and purification of native human lactopontin from human breast milk. The authors also thank Adam Dowle at the Centre of Excellence in Mass Spectrometry, University of York for the MALDI-TOF analysis of glycoform structures.

